# RNA-guided Retargeting of *Sleeping Beauty* Transposition in Human Cells

**DOI:** 10.1101/848309

**Authors:** Adrian Kovač, Csaba Miskey, Michael Menzel, Esther Grueso, Andreas Gogol-Döring, Zoltán Ivics

## Abstract

Two different approaches of genomic modification are currently used for genome engineering and gene therapy: integrating vectors, which can efficiently integrate large transgenes but are unspecific with respect to their integration sites, and nuclease-based approaches, which are highly specific but not efficient at integrating large genetic cargoes. Here we demonstrate biased genome-wide integration of the *Sleeping Beauty* (SB) transposon by combining it with components of the CRISPR/Cas9 system. We provide proof-of-concept that it is possible to influence the target site selection of SB by fusing it to a catalytically inactive Cas9 (dCas9) and by providing a single guide RNA (sgRNA) against the human *Alu* retrotransposon. Enrichment of transposon integrations was dependent on the sgRNA, occurred in a relatively narrow, ∼200 bp window around the targeted sites and displayed an asymmetric pattern with a bias towards sites that are downstream of the sgRNA targets. Our data indicate that the targeting mechanism specified by CRISPR/Cas9 forces integration into genomic regions that are otherwise poor targets for SB transposition. Future modifications of this technology may allow the development of methods for efficient and specific gene insertion for precision genetic engineering.

## INTRODUCTION

The ability to add, remove or modify genes enables researchers to investigate genotype-phenotype relationships in biomedical model systems (functional genomics), to exploit genetic engineering in species of agricultural and industrial interest (biotechnology) and to replace malfunctioning genes or to add functional gene sequences to cells in order to correct diseases at the genetic level (gene therapy).

One option for the insertion of genetic cargo into genomes is the use of integrating vectors. The most widely used integrating genetic vectors were derived from retroviruses, in particular γ-retroviruses and lentiviruses (1). These viruses have the capability of shuttling a transgene into target cells and stably integrating it into the genome, resulting in long-lasting expression. Transposons represent another category of integrating vector. In contrast to retroviruses, transposon-based vectors only consist of a transgene flanked by inverted terminal repeats (ITRs) and a transposase enzyme, the functional equivalent of the retroviral integrase (2). For DNA transposons, the transposase enzymes excise genetic information flanked by the ITRs from the genome or a plasmid and reintegrate it at another position (Fig. 1A). Thus, transposons can be developed as non-viral gene delivery tools (3) that are simpler and cheaper to produce, handle and store than retroviruses (4). The absence of viral proteins may also prevent the immune reactions observed with some viral vectors (5, 6). The *Sleeping Beauty* (SB) transposon is a Class II DNA transposon, whose utility has been demonstrated in pre-clinical [reviewed in (2, 7)] as well as clinical studies [(8, 9) and reviewed in (10)]. It is active across a wide range of cell types (11, 12) and hyperactive variants such as the SB100X transposase catalyze gene transfer in human cells with high efficiency (13).

**Figure 1.**
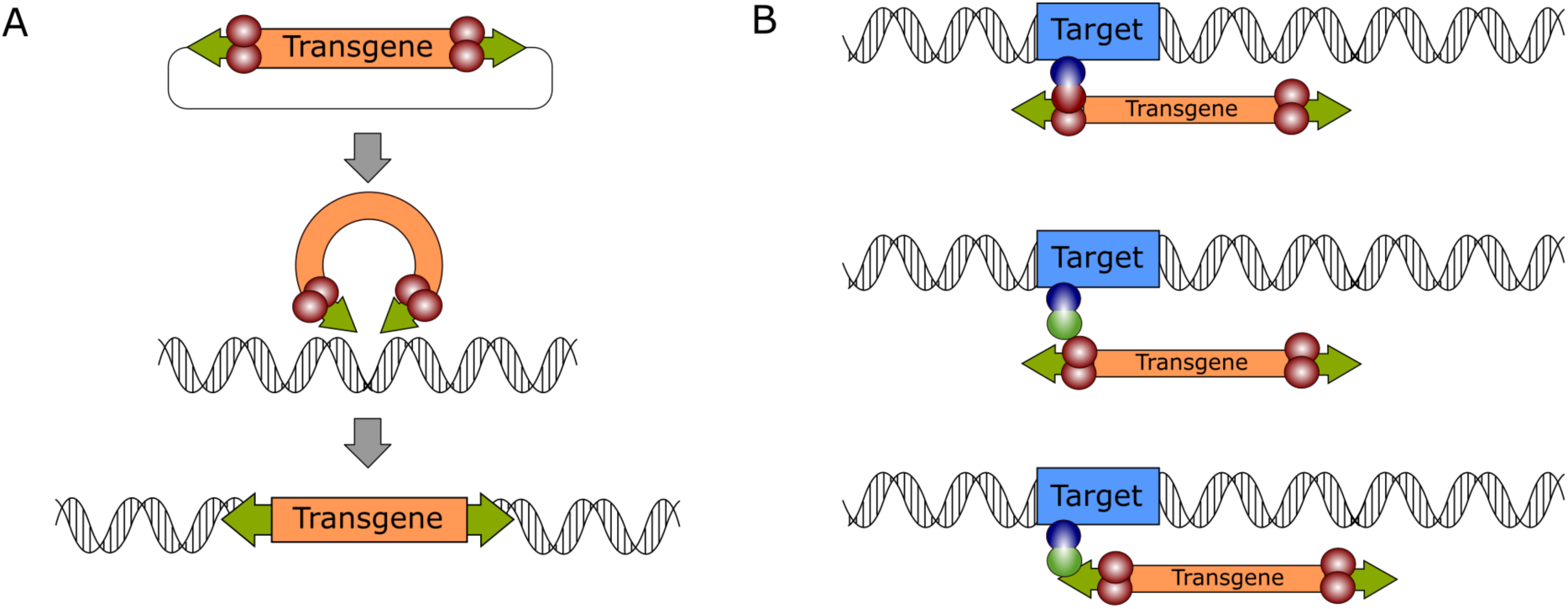
General mechanism of DNA transposition and molecular strategies for targeted gene integration. **(A)** The transpositional mechanism of a DNA transposon in a biotechnological context. The transgene, which is flanked by transposon ITRs (green arrows) is excised from a plasmid by the transposase enzyme (red spheres), which is supplied *in trans.* The genetic cargo is then integrated in the target genome. (**B)** Transposition can be retargeted by foreign factors that can be DNA-binding domains (blue spheres) directly fused to the transposase (top), or to adapter domains (green spheres) that interact either with the transposase (middle) or the transposon DNA (bottom).

The main drawback of integrating vectors is their unspecific or semi-random integration (14). For example, lentiviruses or γ-retroviruses actively target genes or transcriptional start sites (15–19). In contrast, the SB transposon displays a great deal of specificity of insertion at the primary DNA sequence level – almost exclusively integrating into TA dinucleotides (20) – but inserts randomly on a genome-wide scale (21–24). Thus, because all of these vectors can potentially integrate their genetic cargo at a vast number of sites in the genome, the interactions between the transgene and the target genome are difficult to predict. For example, the position of a transgene in the genome can have an effect on the expression of the transgene, endogenous genes or both (25–30). Especially in therapeutic applications, controlled transgene expression levels are important as low expression levels could fail to produce the desired therapeutic effect, while overexpression might in some cases have deleterious effects on the target cell. Perhaps more dramatic are the effects transgenes might have on the genome. Insertion of transgenes can disrupt genomic regulation, either by direct insertional mutagenesis of cellular genes or regulatory elements, or by upregulation of genes in the vicinity of the integration site. In the worst case, this can result in overexpression of a proto-oncogene or disruption of a tumor suppressor gene; both of these outcomes can result in transformation of the cell and tumor formation in the patient.

An alternative technology used in genetic engineering is based on targeted nucleases; the most commonly used nuclease families are zinc finger nucleases (ZFNs) (31), transcription activator-like effector-based nucleases (TALENs) (32) and the CRISPR/Cas system (33). All of these enzymes perform two functions: they have a DNA-binding domain (DBD) that recognizes a specific target sequence and a nuclease domain that cleaves the target DNA once it is bound. While for ZFNs and TALENs target specificity is determined by their amino acid sequence, Cas nucleases need to be supplied with a single guide RNA (sgRNA) that determines their target specificity (34). This makes the CRISPR/Cas system significantly more flexible than other designer nucleases.

The introduction of a double-strand break (DSB) in a target cell is usually repaired by the cell’s DNA repair machinery, either via non-homologous end-joining (NHEJ) or homologous recombination (HR) (35, 36). The NHEJ pathway directly fuses the two DNA ends together. Due to the error-prone nature of this reaction, short insertions or deletions (indels) are often produced. Because this in turn often results in a frame-shift in a coding sequence, this process can be used to effectively knock out genes in target cells. If a DNA template is provided along with the nuclease, a DSB can also be repaired by the HR pathway. This copies the sequence information from the repair template into the target genome, allowing replacement of endogenous sequences or knock-in of completely new genes (37). Thus, knock-in of exogenous sequences into a genetic locus is a cumulative outcome of DNA cleavage by the nuclease and HR by the cell. However, the efficiency of the HR pathway is low compared to the efficiency of the nuclease (38). This bottleneck means that targeted nucleases are highly efficient at knocking out genes (39, 40), but less efficient at inserting DNA (41), particularly when compared to the integrating viral and non-viral vectors mentioned previously. Thus, integrating vectors and nuclease-based approaches to genome engineering have overlapping but distinct advantages and applications: nuclease-based approaches are site-specific and efficient at generating knock-outs, while integrating vectors are unspecific but highly efficient at generating knock-ins.

Based on the features outlined above, it is plausible that the specific advantages of both approaches (designer nucleases and integrating vector systems) could be combined into a single system with the goal of constructing a gene delivery tool, which inserts genetic material into the target cell’s genome with great efficiency and at the same time in a site-specific manner. Indeed, by using DBDs to tether integrating enzymes (retroviral integrases or transposases) to the desired target, one can combine the efficient, DSB-free insertion of genetic cargo with the target specificity of designer nucleases [reviewed in (14)]. In general, two approaches can be used to direct transposon integrations by using a DBD: direct fusions or adapter proteins (14). In the direct fusion approach, a fusion protein of a DBD and the transposase is generated to tether the transposase to the target site (Fig. 1B, top). However, the overall transposase activity of these fusion proteins is often reduced. Alternatively, an adapter protein can be generated by fusing the DBD to a protein domain interacting with the transposon or the transposase (Fig. 1B, middle and bottom). Several transposon systems, notably the SB and the *piggyBac* systems have been successfully targeted towards genomic sites or regions [reviewed in (14)]. The SB transposon system has been successfully targeted to a range of exogenous or endogenous loci in the human genome (42–44). However, a consistent finding across all targeted transposition studies is that while some bias can be introduced, the number of targeted integrations is relatively low when compared to the number of untargeted background integrations (14).

In the studies mentioned above, targeting was achieved with DBDs including ZFs or TALEs, which target a specific sequence determined by their structure. However, for knock-outs, the CRISPR/Cas system is currently the most widely used technology due to its flexibility in design. A catalytically inactive variant of Cas9 called dCas9 (‘dead Cas9’, containing the mutations D10A and H840A), has previously been used to target enzymes including transcriptional activators (45–47), repressors (48, 49), base editors (50, 51) and others (52, 53) to specific target sequences. Using dCas9 as a targeting domain for a transposon could combine this great flexibility with the advantages of integrating vectors. By using the *Hsmar1* human transposon (54), a 15-fold enrichment of transposon insertions into a 600-bp target region was observed in an *in vitro* plasmid-to-plasmid assay employing a dCas9-transposase fusion (55). However, no targeted transposition was detected with this system in bacterial cells. A previous study failed to target the *piggyBac* transposon into the *HPRT* gene with CRISPR/Cas9 components in human cells, even though some targeting was observed with other DBDs (56). However, in a recent study, some integrations were successfully biased to the *CCR5* locus using a dCas9-*piggyBac* fusion (57). Two additional recent studies showed highly specific targeting of bacterial Tn7-like transposons by an RNA-guided mechanism, but only in bacterial cells (58, 59).

Previous studies have established that foreign DBDs specifying binding to both single copy as well as repetitive targets can introduce a bias into SB’s insertion profile, both as direct fusions with the transposase and as fusions to the N57 targeting domain. N57 is an N-terminal fragment of the SB transposase encompassing the N-terminal helix-turn-helix domain of the SB transposase with dual DNA-binding and protein dimerization functions (60). Fusions of N57 with the tetracycline repressor (TetR), the E2C zinc finger domain (61), the ZF-B zinc finger domain and the DBD of the Rep protein of adeno-associated virus (AAV) were previously shown to direct transposition catalyzed by wild-type SB transposase to the *erbB-2* gene, endogenous human L1 retrotransposons and Rep-recognition sequences, respectively (61,42,43,44). Here, we present proof-of-principle evidence that integrations of the SB transposon system can be biased towards loci of choice in the human genome using dCas9 as a targeting domain.

## MATERIALS and METHODS

### Cell culture and transfection

HeLa cells were cultured at 37°C and 5% CO_2_ in DMEM (Gibco) supplemented with 10% (v/v) FCS, 2 mM L-Glutamine (Sigma) and penicillin-streptomycin. For selection, media were supplemented with puromycin (InvivoGen) at 1 µg/ml or 6-thioguanine (Sigma) at 30 mM. Transfections were performed with Lipofectamine 3000 (Invitrogen) according to manufacturer’s instructions.

### Plasmid construction

All sequences of primers and other oligos are listed in **Supplementary Table S1**. dCas9 fusion constructs were generated using pAC2-dual-dCas9VP48-sgExpression (Addgene, #48236) as a starting point. The VP48 activation domain was removed from this vector by digestion with *Fse*I and *Eco*RI. For dCas9-SB100X, the SB100X insert was generated by PCR amplification from a pCMV-SB100X expression plasmid with primers SBfwd_1 (which introduced the first half of the linker sequence) and SBrev_1 (which introduced the *Eco*RI site). The resulting product was PCR amplified using SBfwd_2 and SBrev_1 (SBfwd_2 completed the linker sequence and introduced the *Fse*I site). The generated PCR product was purified, digested with *Eco*RI and *Fse*I and cloned into the dCas9 vector. The dCas9-N57 construct was generated in an analogous manner, replacing primer SBrev_1 with N57rev_1 to generate a shorter insert which included a stop codon in front of the *Eco*RI site. In addition, annealing of phosphorylated oligos stop_top and stop_btm resulted in a short insert containing a stop codon and sticky ends compatible with *Fse*I-and *Eco*RI-digested DNA. Ligation of this oligo into the *Fse*I/*Eco*RI-digested dCas9-VP48 vector resulted in a dCas9 expression plasmid. To generate the N57-dCas9 plasmid, the previously constructed dCas9 expression vector was digested with *Age*I and the N57 sequence was PCR-amplified by two PCRs (using primers SBfwd_3 and N57rev_2, followed by SBfwd_3 and N57rev_3), which introduced a linker and two terminal *Age*I sites. The *Age*I-digested PCR product was ligated into the dCas9 vector, generating a N57-dCas9 expression vector. For Cas9-SB100X and Cas9-N57 constructs, the same cloning strategy was used, using the plasmid pSpCas9(BB)-2A-GFP (Addgene, #113194) as a starting point instead of pAC2-dual-dCas9VP48-sgExpression. Insertion of sgRNAs into Cas9/dCas9-based vectors was performed by digesting the vector backbone with *Bbs*I and inserting gRNA target oligos generated by annealing phosphorylated oligos that included overhangs compatible to the *Bbs*I-digested backbones. For expression, plasmids were transformed into *E. coli* (DH5α or TOP10, Invitrogen) using a standard heat shock protocol, selected on LB agar plates containing ampicillin and clones were cultured in LB medium with ampicillin. Plasmids were isolated using miniprep or midiprep kits (Qiagen or Zymo, respectively).

### *In vitro* Cas9 cleavage assay

For *in vitro* Cas9 cleavage reactions, 1 µg of genomic DNA was incubated with 3 µg of *in vitro* transcribed sgRNAs and 3 µg of purified Cas9 protein in 20µl of 1x NEB3 buffer (New England Biolabs) at 37°C over night. DNA was visualized by agarose gel electrophoresis in a 1% agarose gel. sgRNAs were generated by PCR amplifying the sgRNA sequences with a primer introducing a T7 promoter upstream of the sgRNA and performing *in vitro* transcription using MEGAshortscript™ T7 Transcription Kit (Thermo Fisher). After digestion, fragmented gDNA was purified using a column purification kit (Zymo) and ligated into SmaI-digested pUC19. The plasmids were transformed into *E.coli* DH5α and grown on LB agar supplemented with X-gal. Plasmids from white colonies were isolated and the insert ends were sequenced using primers pUC3 and pUC4. Sanger sequencing was performed by GATC Biotech.

### Western Blot

Protein extracts used for Western Blot were generated by transfecting 5 x 10^6^ HeLa cells with 10 µg of expression vector DNA and lysing cells with RIPA buffer after 48 hours. Lysates were passed through a 23-gauge needle, incubated 30 min on ice, then centrifuged at 10.000 g and 4°C for 10 minutes to remove cell debris. Total protein concentrations were determined via Bradford assay (Pierce™ Coomassie Plus (Bradford) Assay Kit, Thermo Fisher). Proteins were separated by discontinuous SDS-PAGE and transferred onto nitrocellulose membranes (1 hour at 100 V). Membranes were stained with α-SB antibody (1:500, 2 hours) and α-goat-HRP (1:10000, 1 hour) or with α-actin (1:5000, 2 hours) and α-rabbit-HRP (1:10000, 1 hour) for the loading control. Membranes were visualized using ECL™ Prime Western Blotting reagents.

### Transposition assay

Transposition assays were performed by transfecting 10^6^ HeLa cells with 500 ng pT2Bpuro and 10 ng pCMV-SB100X or 20 ng of dCas9-SB100X expression vector. Selection was started 48 hours post-transfection in 10 cm dishes. After two weeks, cells were fixed for two hours with 4 % paraformaldehyde and stained over night with methylene blue. Plates were scanned and colony numbers were automatically determined using ImageJ and the Colony Counter plugin (settings: size > 150 px, circularity > 0.7).

### Cas9 fusion cleavage assay

10^6^ HeLa cells were transfected with 3 µg of Cas9 (without sgRNA), sgHPRT-Cas9, sgHPRT-Cas9-N57or sgHPRT-Cas9-SB100X. Selection with 6-TG was started 72 hours after transfection. Fixing, staining and counting of colonies were performed as detailed in the previous section.

### Electrophoretic mobility shift assay (EMSA)

Nuclear extracts of dCas9-, dCas9-N57-and N57-dCas9-transfected HeLa cells were generated using NE-PER™ Nuclear and Cytoplasmic Extraction Reagents (Thermo Fisher) according to manufacturer’s instructions and total protein concentration was determined by Bradford assay. Similar expression levels between extracts were verified by dot blot using a Cas9 antibody (Cas9 Monoclonal Antibody 7A9, Thermo Fisher). A bacterial extract of N57 was used as a positive control. For the EMSA, a LightShift™ Chemiluminescent EMSA Kit (Thermo Fisher) was used according to manufacturer’s instructions, using ca. 10 µg of total protein (nuclear extracts) or 2.5 µg of total protein (bacterial extract).

### Generation of integration libraries

SB integrations were generated by transfecting 5 x 10^6^ HeLa cells with expression plasmids of either dCas9-SB100X (750 ng) or dCas9-N57 (9 µg). 250 ng of SB100X were co-transfected with dCas9-N57 (but not dCas9-SB100X). All samples were also transfected with 2.5 µg of pTpuroDR3. For each targeting construct, plasmids containing either no sgRNA, the sgRNA HPRT-0 or the sgRNA AluY-1 were used. Puromycin selection was started 48 hours after transfection and cells were cultured for two weeks. Cells were then harvested and genomic DNA was prepared using a DNeasy Blood & Tissue Kit (Qiagen). The protocol and the oligonucleotides for the construction of the insertion libraries have previously been described (62). Briefly, genomic DNA was sonicated to an average length of 600 bp using a Covaris M220 ultrasonicator. Fragmented DNA was subjected to end repair, dA-tailing and linker ligation steps. Transposon-genome junctions were then amplified by nested PCRs using two primer pairs binding to the transposon ITR and the linker, respectively. The PCR products were separated on a 1.5 % ultrapure agarose gel and a size range of 200-500 bp was extracted from the gel. Some of the generated product was cloned and Sanger sequenced for library verification, .before high-throughput sequencing with a NextSeq (Illumina) instrument with single-end 150bp setting.

### Sequencing and bioinformatic analysis

The raw Illumina reads were processed in the R environment (63), as follows: the transposon-specific primer sequences were searched and removed, PCR-specificity was controlled by the presence of transposon end sequences downstream of the primer. After trimming the 5’ of the reads quality trimming was performed if the average phred score values of four consecutive bases dropped below 20. The minimum read-length filter was set to 30. The reads were then mapped to the hg38 human genome assembly using bowtie (64) harnessing the cycling-mapping possibility of TAPDANCE (65). Only single (unambiguous) alignment loci were used for further analysis. Any genomic position was considered valid if supported by at least five independent reads. Since the alignment settings allowed for up to three mismatches independent insertion sites within 5 nucleotides were reduced to the one supported by the highest number of reads. Insertion site logos were calculated and plotted with the SeqLogo package. The frequency of insertions around the sgRNA target sequences were displayed by the genomation package (66). Probability values for nucleosome occupancy in the vicinity of AluY targets and insertion sites were calculated with the algorithm published here (67).

### Statistical analysis

Significance of numerical differences in transposition assay and Cas9 cleavage assays was calculated by performing a two-tailed Student’s t-test using the GraphPad QuickCalcs online tool. All experiments were performed in duplicates or triplicates. We used the Fishers’ exact test for the statistical analyses of the TA-target contents and the frequencies of insertion sites in various genomic intervals.

## RESULTS

### Design and validation of sgRNAs targeting single-copy and repetitive sites in the human genome

Two different targets were chosen for targeting experiments: the *HPRT* gene on the X chromosome and *Alu*Y, an abundant (∼130,000 elements per human genome) and highly conserved family of *Alu* retrotransposons (68). Four sgRNAs were designed to target the *HPRT* gene (Fig. 2A), one of them (sgRNA 0) binding in exon 7 and three (sgRNAs 1-3) in exon 3. Three sgRNAs were designed against *Alu*Y (Fig. 2B), the first two (sgRNAs 1 and 2) against the conserved A-box of the Pol III promoter that drives *Alu* transcription and the third (sgRNA 3) against the A-rich stretch that separates the two monomers in the full-length *Alu* element.

**Figure 2.**
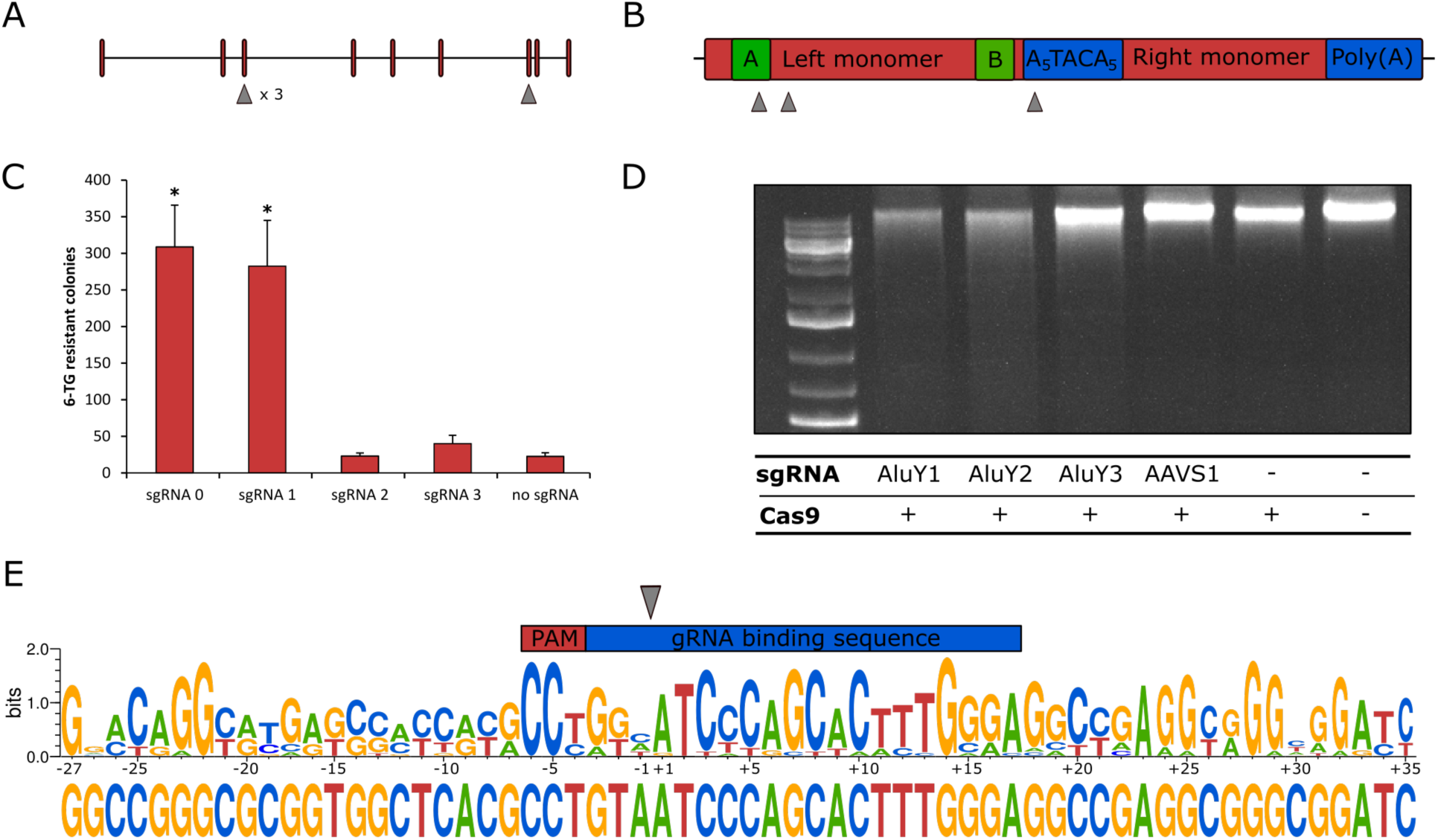
CRISPR/Cas components and their validation for transposon targeting. **(A)** Schematic exon-intron structure of the *HPRT* gene and positions of the sgRNA binding sites. (**B)** Structure of an *Alu* element and relative positions of sgRNA binding sites. (**C)** Number of 6-TG resistant colonies after treatment with Cas9 and *HPRT*-directed sgRNAs. Significance is calculated in comparison to the -sgRNA sample (n=2, biological replicates for all samples, * *p*≤0.05, error bars represent SEM). (**D)** Agarose gel electrophoresis of HEK293T gDNA digested with Cas9 and *Alu*Y-directed sgRNAs. (**E)** Sequence logo generated by aligning sequenced gDNA ends after fragmentation with Cas9 and the *Alu*Y-directed sgRNA (the sequence represents the top strand targeted by the sgRNA). The position of the sgRNA-binding site and PAM is indicated on the top, the cleavage site is marked by the gray arrow. The sequence upstream of the cleavage site is generated from 12 individual sequences, the sequence downstream is generated from 19 individual sequences. The lower sequence represents the *Alu*Y consensus sequence.

The *HPRT*-specific sgRNAs were tested by transfecting human HCT116 cells with a Cas9 expression plasmid and expression plasmids that supply the different *HPRT*-directed sgRNAs. Disruption of the *HPRT* coding sequence by NHEJ was measured by selection with 6-TG, which is lethal to cells in which the *HPRT* gene is intact. Thus, the number of 6-TG-resistant cell colonies obtained in each sample is directly proportional to the extent, to which the *HPRT* coding sequence is mutagenized and functionally inactivated. Two sgRNAs (HPRT-0, HPRT-1) resulted in significantly increased disruption levels (p≤0.05), while the other two (HPRT-2, HPRT-3) failed to increase disruption over the background level obtained in the absence of any sgRNA (Fig. 2C).

The activities of the *Alu*Y-directed sgRNAs were first analyzed by an *in vitro* cleavage assay. Incubation of human genomic DNA (gDNA) with purified Cas9 protein and *in vitro* transcribed sgRNAs showed detectable fragmentation of gDNA for sgRNAs AluY-1 and AluY-2 (Fig. 2D). gDNA digested with Cas9 and sgRNA AluY-1 was purified, cloned into a plasmid vector and the sequences of the plasmid-genomic DNA junctions were determined. 12 of 32 sequenced genomic junctions could be mapped to the *Alu*Y sequence upstream of the cleavage site and 19 could be mapped to the sequence immediately downstream (we use the term upstream to refer to the direction of the left monomer). A consensus sequence generated by aligning the 12 or 19 sequences shows significant similarity to the *Alu*Y consensus sequence (Fig. 2E), demonstrating that the DNA fragmentation was indeed the result of Cas9-mediated cleavage. The sequence composition also shows that mismatches within the sgRNA binding sequence are tolerated to some extent, while other segments including the conserved GG dinucleotide of the NGG PAM motif do not show any sequence variation (Fig. 2E). These data establish efficient sgRNAs against both single-copy and multi-copy targets in the human genome as measured by the activity of Cas9 at these targets in *in vitro* and in cell-based cleavage assays.

### Generation of Cas9 fusion constructs and their functional validation

Three different targeting constructs were generated to test both the direct fusion and the adapter protein approaches described above. For the direct fusion, the entire coding sequence of SB100X, a hyperactive version of the SB transposase (13), was inserted at the C-terminus of the dCas9 sequence (Fig. 3A, top).

**Figure 3.**
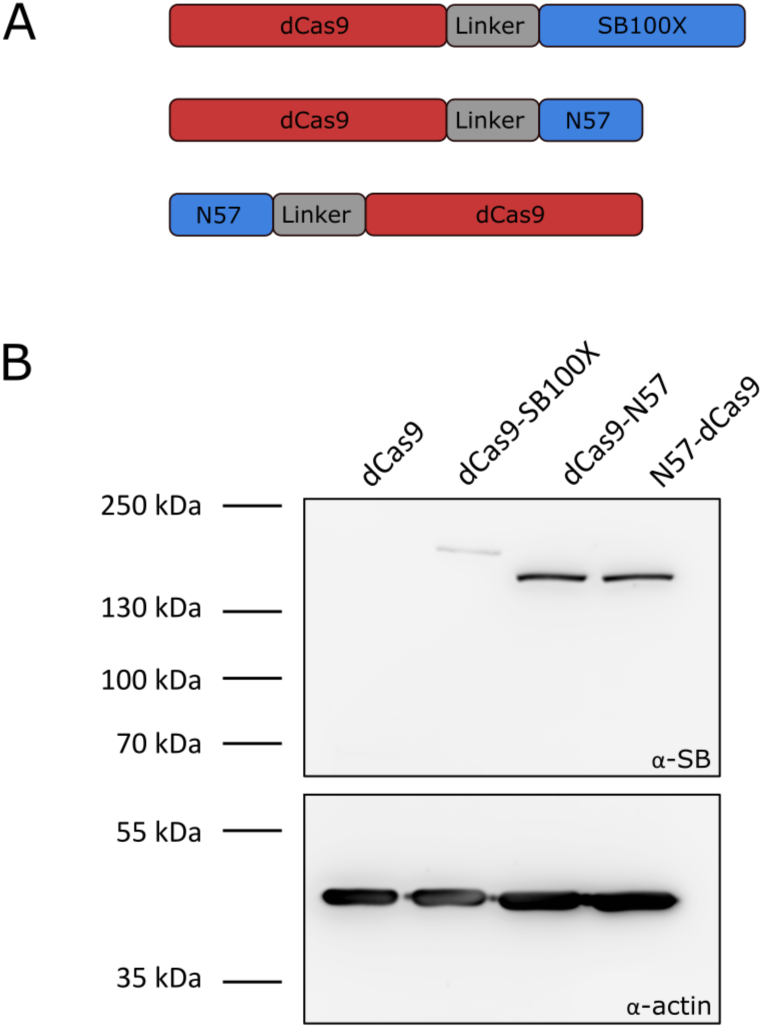
Transposase-derived targeting factors. **(A)** Schematic representation of the targeting constructs. (**B)** Western blot of proteins expressed by the targeting constructs. The top half of the membrane was treated with α-SB antibody, the bottom half was treated with α-actin as a loading control. Expected sizes were 202.5 kDa for dCas9-SB100X and 169.7 kDa for dCas9-N57 and N57-dCas9.

We only made an N-terminal SB fusion, because C-terminal tagging of the transposase enzyme completely abolish its activity (69,44,70). For adapter proteins, the N57 targeting domain, which interacts both with the SB transposase as well as with the transposon DNA and could therefore recruit either or both components of the SB system to the target site, was inserted at the N-terminus as well as at the C-terminus of dCas9 (Fig. 3A, middle and bottom, respectively). A flexible linker KLGGGAPAVGGGPK (71) that was previously validated in the context of SB transposase fusions to ZF (42) and to Rep (43) DBDs was introduced between dCas9 and the full-length SB100X transposase or the N57 targeting domain (Fig. 3A). All three protein fusions were cloned into an all-in-one expression plasmids that allow co-expression of the dCas9-based targeting factors with sgRNAs.

Western blots using an antibody against the SB transposase verified the integrity and the expression of the fusion proteins (Fig. 3B). In order to verify that the dCas9-SB100X direct fusion retained sufficient transpositional activity we measured its efficiency at integrating a puromycin-marked transposon into HeLa cells and compared its activity to the unfused SB100X transposase (Fig. 4A). We found that the fusion construct dCas9-SB100X was approximately 30 % as active as the unfused SB100X. To verify that N57 retains its DNA-binding activity in the context of the dCas9 fusions we performed an EMSA experiment using a short double-stranded oligonucleotide corresponding to the N57 binding sequence in the SB transposon (Fig. 4B). Binding could be detected for the dCas9-N57 fusion, but not for N57-dCas9. For this reason, the N57-dCas9 construct was excluded from the subsequent experiments. The DNA-binding ability of the dCas9 domain in the fusion constructs was tested by generating alternative constructs that included catalytically active Cas9 domains as a proxy, but were otherwise identical. The activity of these fusion constructs was determined by measuring the disruption frequency of the *HPRT* gene by selection with 6-TG (Fig. 4C) as described above. For both Cas9-SB100X as well as Cas9-N57, the cleavage efficiency was determined to be ∼30 % of unfused Cas9 in the presence of sgRNA HPRT-0. Since target DNA binding is a prerequisite for Cas9 cleavage, and the overall structure of Cas9 is very similar to dCas9, we assumed that the dCas9 domain retains significant DNA-binding ability in both fusion constructs.

**Figure 4.**
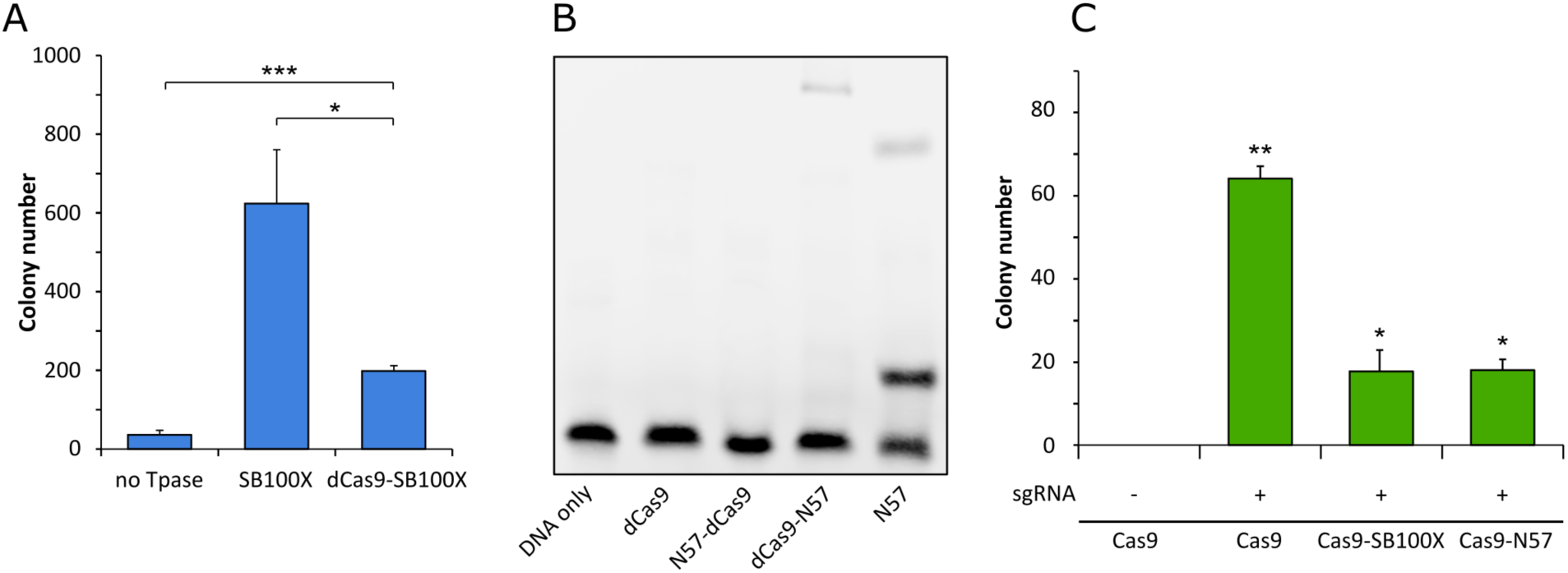
Functional testing of dCas9 fusions. **(A)** Numbers of puromycin-resistant colonies in the transposition assay. The dCas9-SB100X fusion protein catalyzes ∼30% as many integration events as unfused SB100X transposase (n=3, biological replicates, * *p*≤0.05, *** *p*≤0.001, error bars represent SEM). (**B)** EMSA with dCas9-N57 fusion proteins. dCas9 serves as negative control, N57 as positive control. Binding can be detected for dCas9-N57, but not for N57-dCas9. (**C)** Numbers of 6-TG resistant colonies after Cas9 cleavage assay. No disruption of the *HPRT* gene, as measured by 6-TG resistance, can be detected without the addition of the HPRT sgRNA. In the presence of a HPRT sgRNA, all Cas9 constructs cause significant disruption of the *HPRT* gene (n=2, biological replicates, * *p*≤0.05, ** *p*≤0.01, error bars represent SEM).

Collectively, these data establish that our dCas9 fusion proteins i) are active in binding to the target DNA in the presence of sgRNA; ii) they retain transposition activity (for the fusion with the full-length SB100X transposase); and iii) they can bind to the transposon DNA (for the fusion with the C-terminal N57 targeting domain), which constitute the minimal requirements for targeted transposition in the human genome.

### RNA-guided *Sleeping Beauty* transposition in the human genome

Having established functionality of our multi-component transposon targeting system, we next analyzed the genome-wide patterns of transposon integrations catalyzed by the different constructs. Transposition reactions were performed in human HeLa cells with dCas9-SB100X or dCas9-N57 + SB100X. For both constructs, the reactions were complemented either with sgRNA HPRT-0 or sgRNA AluY-1 or with no sgRNA. Integration libraries consisting of PCR-amplified transposon-genome junctions were generated and subjected to high-throughput sequencing. Recovered reads were aligned to the human genome (hg38 assembly) to generate lists of insertion sites. In order to quantify the targeting effects, we defined targeting windows of increasing lengths around the sgRNA binding sites (Fig. 5A). The number of insertions in each targeting window (Fig. 5B) was converted into a fraction of overall insertions (Fig. 5C) and these ratios were compared to those obtained with the negative control (same targeting construct without sgRNA) (Fig. 5D).

**Figure 5.**
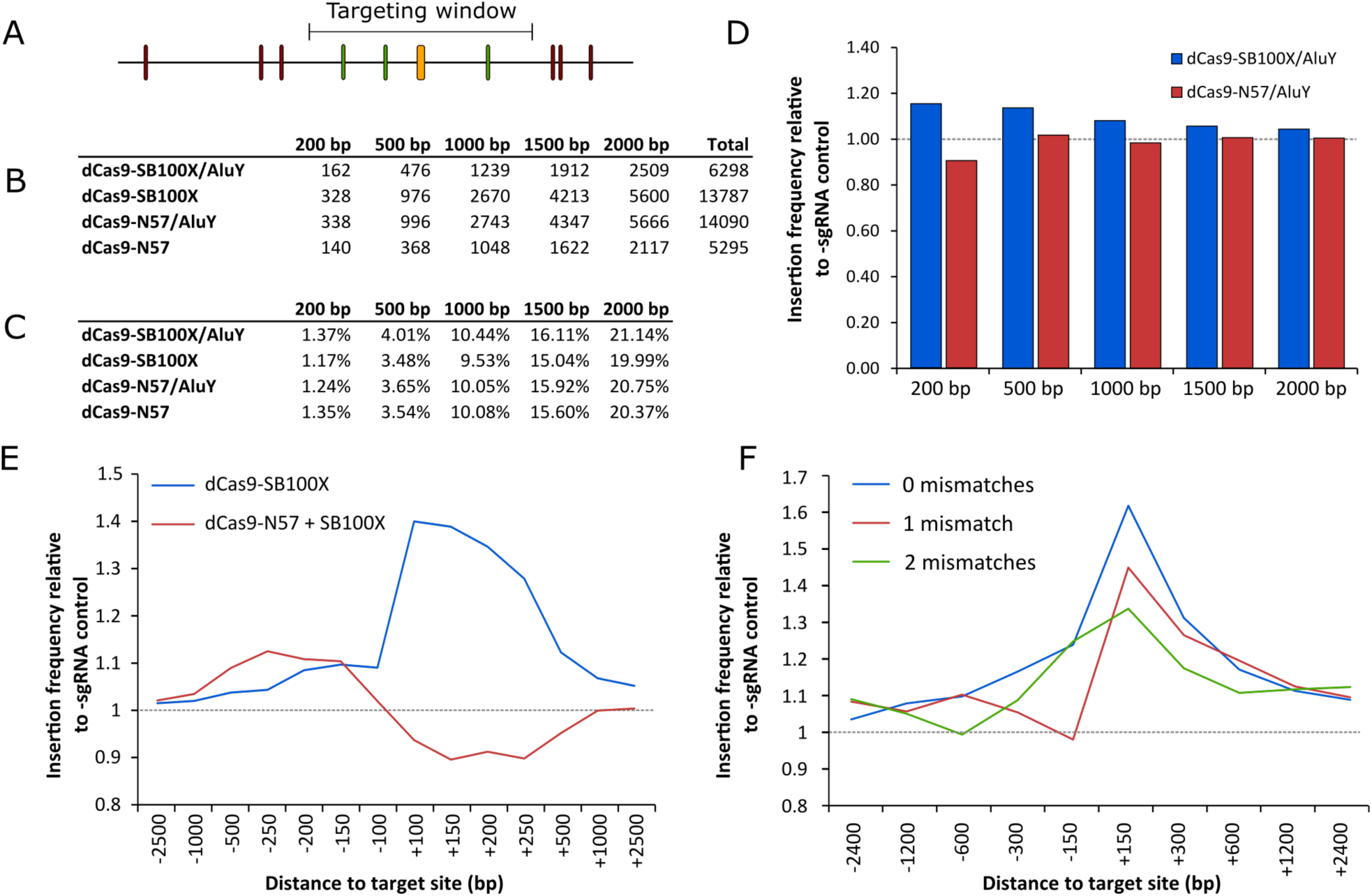
RNA-guided Sleeping Beauty transposition in human cells. **(A)** Schematic representation of the analysis of SB retargeting. Targeting windows are defined as DNA extending a certain number of base pairs upstream or downstream of the sgRNA target sites (yellow – sgRNA target, green – ‘hit’ insertion, red – ‘miss’ insertion). (**B)** Numbers of integrations recovered from windows of different sizes along with the total numbers of integrations in the respective libraries. (**C)** Percentages of insertions that the insertion counts in Figure 5B represent. (**D)** Relative insertion frequencies in windows of various sizes around the targeted sites. The strongest enrichment in the dCas9-SB100X sample occurs in 200-bp windows. Insertion frequencies are normalized with the respective -sgRNA sample. The windows are cumulative, i.e. the 500-bp window also includes insertions from the 200-bp window. (**E)** Relative insertion frequencies in windows of various sizes, upstream and downstream of the target sites. The strongest enrichment in the dCas9-SB100X sample occurs upstream of the sgRNA target site. Insertion frequencies are relative to the respective -sgRNA sample. (**F)** The effect of the number of mismatches on the targeting efficiency of dCas9-SB100X. Relative insertion frequencies of the dCas9-SB100X sample around perfectly matched target sites as well as sites with 1 or 2 mismatches.

For the *HPRT* locus, no insertion was recovered within 5 kb in either direction from the sgRNA binding site. We then included sites with up to three mismatches to the sgRNA target sequence in the target list and looked for enrichment in targeting windows around these 1259 sites. We detected a single insertion within 500 bp of a site with three mismatches to the HPRT sgRNA in the sgHPRT-dCas9-N57 + SB100X sample. While this represents a four-fold higher fraction of total insertions of this library compared to the control (0.12% compared to 0.03%), it is not statistically significant due to the low number of insertions involved.

The sgRNA AluY-1 has a total of 299339 target sites in the human genome (hg38) (the number of sites exceeds the number of *Alu*Y elements due to high conservation, and therefore presence in other *Alu* subfamilies). We found that for dCas9-N57 + SB100X the fractions of insertions falling into the mapping windows were practically identical in the presence and absence of the sgRNA, indicating that targeting with the adapter protein was unsuccessful (Fig. 5D). However, we detected a 15% increase in the frequency of insertions within the 200-bp-wide window around the sgAluY-1 target sites for the Cas9-SB100X fusion in the presence of the sgRNA (1.37 % *vs*. 1.17 %). Enrichment of insertions in the presence of sgRNA was the highest close to the target site (within a 200-bp window) and gradually leveled off in the larger analysis windows (Fig. 5D), indicating a local effect consistent with the proposed targeting mechanism (Fig. 1B), which presumes tethering the transpositional complex in close vicinity of the targeted loci. To investigate potential upstream and downstream biases, we counted insertions in up-and downstream windows independently, and found that the enrichment occurred mainly downstream of the target sites (Fig. 5E), within the *Alu*Y element. We also detected enrichment near target loci similar to the target site (up to 2 mismatches) (Fig. 5F). This result is in agreement with the finding that the specificity of dCas9 binding is lower than that of Cas9 cleavage (72). As expected, targeting efficiency decreased with the number of mismatches. We detected statistically significant enrichments in the insertion frequencies in windows of 300 bps embracing target sites of 0 and 1 mismatch (∼1.6-fold enrichment, *p*=0.022 and ∼1.4-fold enrichment, *p*=0.003), respectively.

Intriguingly, plotting the overall insertion frequencies around the target sites revealed that the SB insertion machinery generally disfavors loci in the immediate vicinity of the sgRNA binding sequences (Fig. 6A). These results together with the asymmetric pattern of integrations next to the target sites prompted us to investigate properties of the genomic loci around the sgRNA target sites. To our surprise, we found that the TA dinucleotide frequency in the targeted region is in fact lower than in the neighboring segments (Fig. 6B), thus targeting occurs into *per se* disfavored DNA. Since the nucleotide composition of the targeted regions is remarkably different from that of the neighboring sequences and given that nucleosome positioning in the genome is primarily driven by sequence (67), we next investigated nucleosome occupancy of the target DNA. Nucleosome occupancy was predicted in 2-kb windows on 20000 random target sequences and on all the insertion sites of the non-targeted condition. This analysis recapitulated our previous finding showing that SB disfavors integrating into DNA present in nucleosomes (73). Additionally, in agreement with previous findings of others (74, 75), we found that these *Alu*Y sequences are conserved regions for nucleosome formation (Fig. 6C). These results can explain the overall drop in insertion frequency of SB into these regions. Along this line, we next set out to investigate the target nucleotides of the transposons in the targeted segments. SB inserts predominantly into TA target sites. However, when comparing the frequency of insertions of the fusion construct into non-TA sequences within *vs*. outside the targeted 300-nucleotide windows, we detected an increase in the number of non-TA targeted events 5.35% vs. 3.03% (*p*=0.023) (**Supplementary Fig. S1**). In sum, the data above establish sgRNA-dependent enrichment of SB transposon integrations displaying an asymmetric distribution around multicopy genomic target sites in the human genome.

**Figure 6.**
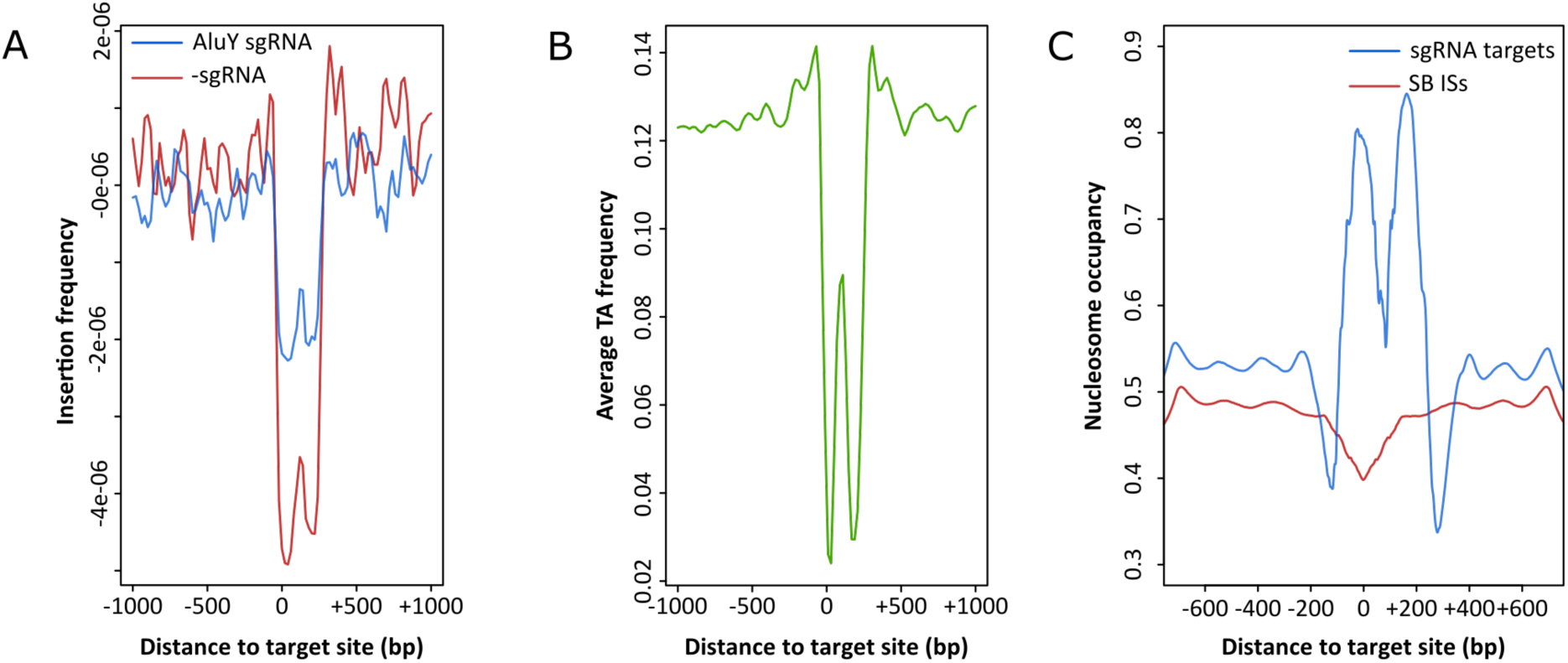
Analysis of targeted chromosomal regions. (**A**) Insertion frequencies of the targeted (blue) and non-targeted (red) dataset show that enrichment occurs within a 300-bp window upstream of the sgRNA target sites, which is generally disfavored for SB integration. (**B**) Reduced average TA di-nucleotide frequency within the targeted 300-bp window. (**C**) Computationally predicted nucleosome occupancy around the sites targeted by the *Alu*Y sgRNA (blue) and around untargeted SB insertion sites (red).

## DISCUSSION

We demonstrate in this study that the insertion pattern of the SB transposase can be influenced by fusion to dCas9 as a RNA-guided targeting domain in human cells, and as a result be biased towards sites specified by a sgRNA that targets a sequence in the *Alu*Y repetitive element. We consider it likely that the observed enrichment of insertions next to sgRNA-targeted sites is an underestimate of the true efficiency of transposon targeting in our experiments, because our PCR procedure followed by next generation sequencing and bioinformatic analysis cannot detect independent targeting events that had occurred at the same TA dinucleotide in the human genome.

Our data indicate a relatively narrow targeting window around the site specified by the sgRNA. This observation is consistent with physical docking of the transpositional complex at the targeted site, and suggests that binding of dCas9 to its target sequence and integration by the SB transposase occur within a short timeframe. We further detect an asymmetric distribution of insertions around the target site. Asymmetric distributions have been found in several previous targeting studies, for example a study using the ISY100 transposon (which, like SB, is a member of the Tc1/*mariner* transposon superfamily) in combination with the ZF domain Zif268 in *E. coli* (76) or *in vitro* experiments with dCas9/Hsmar1 fusions (55). Enrichment mainly occurring downstream of the sgRNA target site was somewhat surprising, as domains fused to the C-terminus of Cas9 are expected to be localized closer to the 5’-end of the target strand (77), or upstream of the sgRNA binding site. The fact that SB100X is connected with dCas9 by a relatively long, flexible linker could explain why enrichment can occur on the other side of the sgRNA binding site, but it does not explain why enrichment on the ‘far side’ seems to be more efficient. Against expectations, we find that the window, in which the highest enrichment occurs, is a disfavored target for SB transposition (Fig. 6A), likely because it is TA-poor (Fig. 6B) – the *Alu*Y consensus sequence has a GC content of 63% (78) – and nucleosomal (Fig. 6C). It is possible that the targeting effect in this window is more pronounced than on the other side of the target site because there are fewer background insertions obscuring the effect. It should also be noted that comparison of sequence logos of insertion sites showed that a higher fraction of insertions occur at non-canonical (i.e. non-TA) sites within the 300-bp window than outside of it (**Supplementary Fig. S1**). The effect of increased non-canonical integration in a targeting context was also observed in another recent transposon retargeting study (57), and might be a further indication that targeting forces a fraction of insertions into non-optimal sites that would likely not have been selected without the constraints imposed by the targeting construct.

Unlike in our earlier studies establishing biased transposon integration by the N57 targeting peptide fused to various DBDs (44,42,43), our dCas9-N57 fusion apparently did not exert a measurable effect on the genome-wide distribution of SB transposon insertions (Fig. 5). Because Cas9-N57 is active in cleavage (Fig. 4C) and dCas9-N57 is active in binding to transposon DNA (Fig. 4B), this result was somewhat unexpected. We speculate that addition of a large protein (dCas9 is 158 kDa) to the N-terminus of a relatively small polypeptide of 57 amino acids masks its function to some extent. Indeed, the TetR, ZF-B protein and Rep DBD that were used previously with success in conjunction with N57 are all far smaller than dCas9. The binding activity of N57 to transposon DNA, though detectable by EMSA, may have been too weak to effectively recruit the components of the SB system to the target site.

Our data reveal some of the important areas where refined molecular strategies as well as reagents may yield higher targeting efficiencies. First, the difficulty of targeting to a single location, in this case the *HPRT* gene, might be associated with characteristics of the target itself or an indication that the system is not specific enough to target a single-copy site in general. The fact that an integration library consisting of 21646 independent SB integrations generated by unfused SB100X without any targeting factor also did not contain any integrations within 50 kb of the *HPRT* target sequence (data not shown) might indicate that the *HPRT* gene is simply a poor target for SB integrations. It should be noted that a previous attempt to target the *piggyBac* transposase to the *HPRT* gene with CRISPR/Cas components also failed, even though targeting with other DBDs (ZFs and TALEs) was successful (56). Poor targeting of a single-copy chromosomal region is reminiscent of our previous findings with engineered Rep proteins (43). Both Rep/SB and Rep/N57 fusions were able to enrich SB transposon integrations in the vicinity of genomic Rep binding sites, yet they failed to target integration into the *AAVS1* locus, the canonical integration site of AAV (43). Thus, selection of an appropriate target site appears to be of paramount importance. The minimal requirements for such sites are accessibility by the transpositional complex and the presence of TA dinucleotides to support SB transposition; in fact, SB was reported to prefer insertion into TA-rich DNA in general (79). The importance of DNA composition in the vicinity of targeted sites was also highlighted in the context of targeted *piggyBac* transposition in human cells (80). Namely, biased transposition was only observed with engineered loci that contained numerous TTAA sites (the target site of *piggyBac* transposons) in the flanking regions of a DNA sequence bound by a ZF protein. An alternative, empirical approach, where careful choice of the targeted chromosomal region may increase targeting efficiencies would be to select sites where clusters of SB insertions (transposition “hot spots”) were identified without a targeting factor. Since these sites are clearly receptive to SB insertions, targeting might be more efficient there. Collectively, these considerations should assist in the design of target-selected gene insertion systems with enhanced efficiency and specificity.

The results presented here, as well as the results of previous targeting studies (14,56,57), indicate that the main obstacle to targeted transposition is the low ratio of targeted to non-targeted insertions. This is likely due to the fact that, in contrast to site-specific nucleases where sequence-specific DNA cleavage is dependent on heterodimerization of *Fok*I endonuclease domain monomers (81), or to Cas9, where DNA cleavage is dependent on a conformational change induced by DNA binding (72), the transposition reaction is not dependent on site-specific target DNA binding. The transposase component, whether as part of a fusion protein or supplied in addition to an adapter protein, is capable of catalyzing integrations without the DBD binding to its target. Thus, any attempt to target specific sites faces an overwhelming excess of non-specific competitor DNA, to which the transposase can freely bind. This non-specific binding of the transposase to human chromosomal DNA competes with specific binding to a desired target sequence, thereby limiting the probabilities of targeted transposition events. This problem might be mitigated by engineering of the transposase to reduce its unspecific DNA affinity. As SB transposase molecules have a positively charged surface (82), they readily bind to DNA regardless of sequence. Decreasing the surface charge of the transposase would likely result in reduced overall activity, but it might make the transposition reaction more dependent on binding to the target DNA by the associated DBD. The ultimate goal would be the design of transposase mutants deficient in target DNA binding but proficient in catalysis. A similar approach was previously applied to *piggyBac* transposase mutants deficient in transposon integration. Although fusion of a ZF DBD restored integration in that study, enrichment of insertion near target sites specified by the DBD was not seen (83). Another simple modification that could potentially result in better targeting is temporal control of the system. In its current form, all components of the system are supplied to the cell at the same time. It might be possible to increase targeting efficiency by supplying the targeting factor first and the transposon at a later point to provide the targeting factors with more time to bind to their target sites.

In conclusion, this study shows that targeting SB transposon integrations towards specific sites in the human genome by an RNA-guided mechanism is possible. This is the first time this has been demonstrated for the SB system and the first time RNA-guided transposition was demonstrated by analyzing the overall distribution of insertion sites on a genome-wide scale. If the current limitations of the system can be addressed, this technology might be attractive for a range of applications including therapeutic cell engineering. Gene targeting by HR is limited in non-dividing cells because HR is generally active in late S and G2 phases of the cell cycle (84). Therefore, post-mitotic cells cannot be edited in this manner (85, 86). Newer gene editing technologies that do not rely on HR, like prime editing (87), usually have a size limitation for insertions that precludes using them to insert entire genes. In contrast, SB transposition is not limited to dividing cells (88) and can transfer genes over 100 kb in size (89). Another drawback of methods relying on generating DSBs is the relative unpredictability of the outcome of editing. As described above, different repair pathways can result in different outcomes at the site of a DSB. Attempts to insert a genetic sequence using HR can also result in the formation of indels or even complex genomic rearrangements (90). In contrast to DSB generation followed by HR, insertion by integrating vectors including transposons occurs as a concerted transesterification reaction (91, 92), avoiding the problems associated with free DNA ends.

## Supplementary data

Supplementary Information (Supplementary Table S1 and Supplementary Figure S1).

## Acknowledgement

We thank T. Diem for technical support.

## Conflict of interest

Z. I. is co-inventor on patents relating to targeted gene insertion.

