## Supplementary Table S1 and Supplementary Figure S1 for "RNA-guided Retargeting of *Sleeping Beauty* Transposition in Human Cells"

**Supplementary Table 1**. Sequences of all used DNA oligos.

| | Oligo sequence |
|---|---|
| **PCR primers** | |
| **SBfwd_1** | ACCTGCTGTGGGCGGAGGCCCTAAGATGGGAAAATCAAAAGAAATCAGCCAAGAC |
| **SBfwd_2** | AATCGGCCGGCCAAACTGGGCGGAGGCGCACCTGCTGTGGGCGGAG |
| **SBfwd_3** | AATCACCGGTACCATGGGAAAATCAAAAGAAATCAGCCAAGAC |
| **SBrev_1** | AATCGAATTCTAGTATTTGGTAGCATTGCCTTTAAATTGTTTAACTT |
| **N57rev** | AATCGAATTCTAGCGGTATGACGGCTGCGTG |
| **N57rev_2** | CAGGTGCGCCTCCGCCCAGTTTGCGGTATGACGGCTGCG |
| **N57rev_3** | AATCACCGGTCTTAGGGCCTCCGCCCACAGCAGGTGCGCCTCCGC |
| **pUC3** | CGATTAAGTTGGGTAACGCCAGGG |
| **pUC4** | GCTGGCACGACAGGTTTCCCG |
| **Other oligos** | |
| **Stop_top** | [Phos]CCTGAG |
| **Stop_btm** | [Phos]AATTCTCAGGCCGG |
| **AluY-1 top** | [Phos]CACCTCCCAAAGTGCTGGGATTAC |
| **AluY-1 bottom** | [Phos]AAACGTAATCCCAGCACTTTGGGA |
| **AluY-2 top** | [Phos]CACCGCCTGTAATCCCAGCACTTT |
| **AluY-2 bottom** | [Phos]AAACAAAGTGCTGGGATTACAGGC |
| **AluY-3 top** | [Phos]CACCTTTTGTATTTTTAGTAGAGA |
| **AluY-3 bottom** | [Phos]AAACTCTCTACTAAAAATACAAAA |
| **HPRT-0 top** | [Phos]CACCGAAGTAATTCACTTACAGTC |
| **HPRT-0 bottom** | [Phos]AAACGACTGTAAGTGAATTACTTC |
| **HPRT-1 top** | [Phos]CACCTCTTGCTCGAGATGTGATGA |
| **HPRT-1 bottom** | [Phos]AAACTCATCACATCTCGAGCAAGA |
| **HPRT-2 top** | [Phos]CACCTAAATTCTTTGCTGACCTGC |
| **HPRT-2 bottom** | [Phos]AAACGCAGGTCAGCAAAGAATTTA |
| **HPRT-3 top** | [Phos]CACCCTGATAAAATCTACAGTCAT |
| **HPRT-3 bottom** | [Phos]AAACATGACTGTAGATTTTATCAG |
| **N57 EMSA top** | TACAGTTGAAGTCGGAAGTTTACATACACTTAAG |
| **N57 EMSA bottom** | CTTAAGTGTATGTAAACTTCCGACTTCAACTGTA |

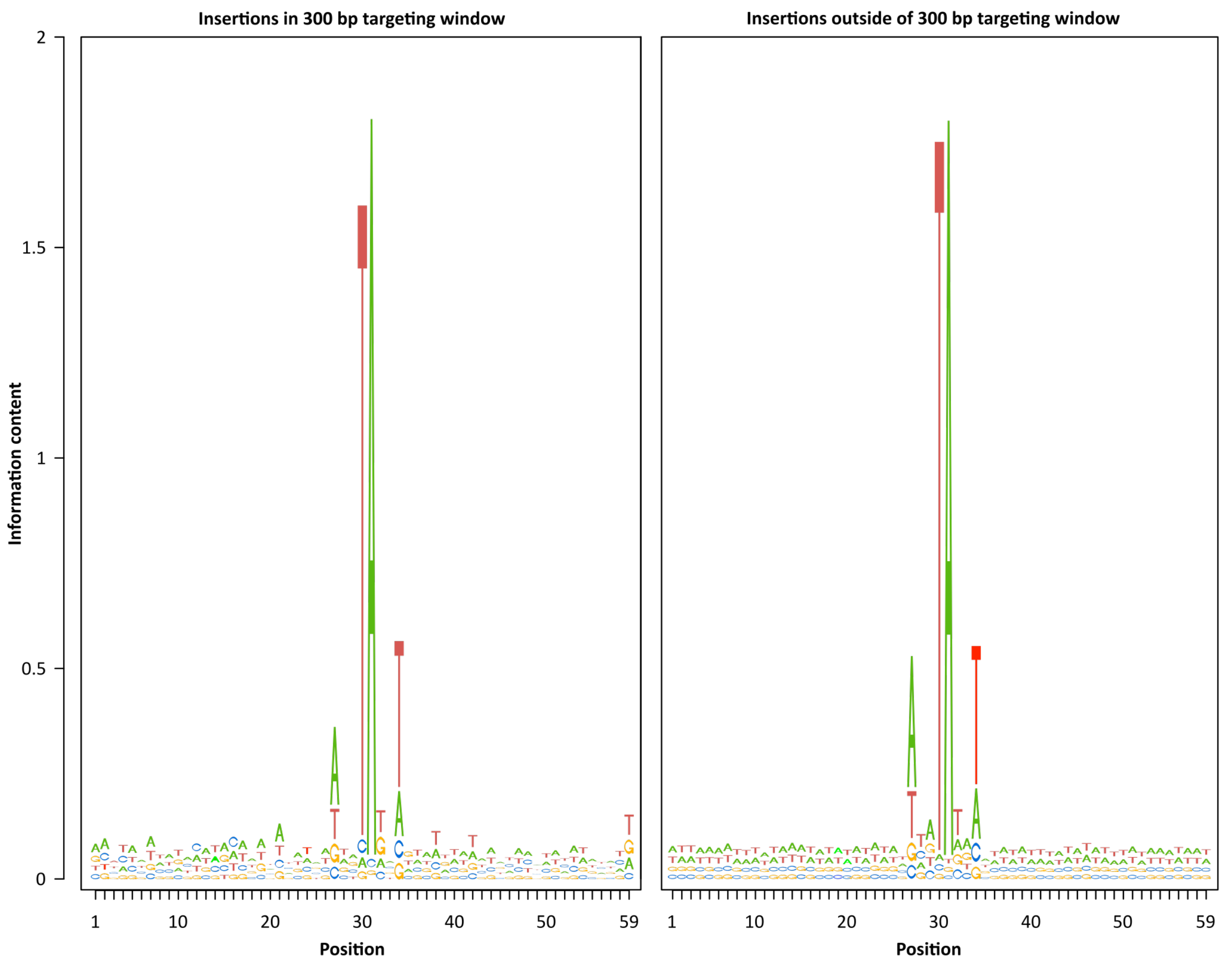


**Supplementary Figure S1**. Sequence logos generated from sequences around insertion sites catalyzed by dCas9-SB100X within the 300-bp targeting window (left) and outside of the window (right). The frequency of insertion into non-TA sites in the targeting window is increased (5.35% vs. 3.03%, $p$=0.023). The most common non-TA dinucleotide into which insertions occurred was CA.
